# Precise Timing Matters: Modulating Human Memory by Synchronizing Hippocampal Stimulation to Saccadic Event Related Potentials

**DOI:** 10.1101/2024.04.09.588460

**Authors:** C.N Katz, K. Patel, A.G. Schjetnan, V. Barkley, David Groppe, J Zariffa, K.D. Duncan, T.A Valiante

**Affiliations:** Krembil Research Institute, Toronto Western Hospital (TWH), Toronto, ON, Canada; Institute of Biomedical Engineering, University of Toronto, Toronto, ON, Canada; Faculty of Medicine, University of Calgary, Calgary, AB, Canada; Center of Advancing Neurotechnological Innovation to Application (CRANIA), University Health Network and University of Toronto, Toronto, ON, Canada; Faculty of Medicine, University of Toronto, Toronto, ON, Canada; Persyst Development Corporation, Solana Beach, CA, United States of America; Edward S. Rogers Sr. Department of Electrical and Computer Engineering, University of Toronto, Toronto, ON, Canada; Rehabilitation Sciences Institute, University of Toronto; KITE Research Institute, Toronto Rehabilitation Institute - University Health Network; Department of Psychology, University of Toronto, Toronto, ON, Canada; Division of Neurosurgery, Department of Surgery, University of Toronto, Toronto, ON, Canada; Institute of Medical Sciences, University of Toronto, Toronto, ON, Canada; Max Planck - University of Toronto Center for Neural Science and Technology; University of Toronto, Toronto, ON, Canada

## Abstract

Episodic memory, the ability to record and relive experiences, is intricately connected to visual exploration in most humans. This study explores the possibility that eye movements create physiological states relevant for memory, analogous to those associated with hippocampal theta. Previous work has demonstrated that saccadic eye movements, which occur roughly at theta frequency, elicit hippocampal event-related potentials (ERPs). Building on the Separate Phases of Encoding and Retrieval (SPEAR) model, we asked if the peaks and troughs of this saccadic ERP are differentially important for memory formation. Specifically, we applied saccade-contingent hippocampal electrical stimulation at estimated ERP peaks and troughs while individuals with epilepsy visually explored natural scenes across 59 sessions. We subsequently assessed their recognition memory for scenes and their recall of associated targets. Results indicate that memory is robust when stimulation precisely targets the peak or trough, contrasting with impairments observed with random stimulation. Moreover, memory impairment is prominent when stimulation is applied within 100 ms of saccade initiation, a time that reflects high medial temporal lobe inhibition. Our findings suggest that the hippocampus rapidly evolves through memory-relevant states following each eye movement, while also challenging the assumption that human saccadic ERP peaks and troughs mirror the encoding and retrieval phases of theta rhythms studied in rodents. The study sheds light on the dynamic interplay between eye movements, hippocampal activity, and memory formation, offering theoretical insights with potential applications for memory modulation in neurological disorders.

**Significance Statement:** Why do eye-movements enhance memory formation? Here, we causally tested if eye-movements initiate short-lived states critical for memory formation within the hippocampus, a brain region known to support memory. We built a system that could precisely apply hippocampal electrical stimulation at key moments after eye-movements to test how the timing of this stimulation influenced people’s ability to form memories. We found that stimulation was particularly disruptive to memory formation when applied within 100 ms of initiating an eye movement. By contrast, memory was robust to stimulation precisely timed to the peak and trough of hippocampal eye-movement responses. We interpret this temporal evolution of memory-relevant states within a prominent model of how theta phases relate to rodent memory.

## Introduction

Since most humans primarily explore their environments visually, it seems intuitive that our eye movements are linked to what we remember (Hannula et al., 2010; Meister and Buffalo, 2016; Ryan et al., 2020). For instance, people better recognize images they freely view (Henderson et al., 2005), even when restricted eye movements are yoked to others’ (Chan et al., 2011). Additionally, the number of eye movements produced during learning is correlated with subsequent memory (Kafkas and Montaldi, 2011; Molitor et al., 2014; Olsen et al., 2016; Fehlmann et al., 2020). The critical question – is why? An evocative possibility is that eye-movements produce physiological states conducive to encoding in the mesial temporal lobe (MTL). Specifically, saccadic eye movements generate event-related potentials (ERP) in the primate MTL (Sobotka and Ringo, 1997; Hoffman et al., 2013; Jutras et al., 2013; Andrillon et al., 2015; Katz et al., 2020). These ERPs reflect heightened periods of functional connectivity (Sobotka et al., 2002) and are dominated by activity in the theta range (3-8 Hz) (Hoffman et al., 2013; Jutras et al., 2013; Katz et al., 2020)—a rhythm linked to memory across species (Buzsáki, 2005; Herweg et al., 2019). Here we causally assess whether the saccadic ERP contains privileged states for human memory.

Our approach was motivated by the SPEAR (<u>S</u>eparate <u>P</u>hases of <u>E</u>ncoding <u>a</u>nd <u>R</u>etrieval) model, which proposes the hippocampus encodes memories during fissure recorded theta’s troughs and retrieves them during the peaks^1^ (Hasselmo et al., 2002; Hasselmo and Stern, 2014). This “gating” function (Womelsdorf et al., 2014) explains why electrical stimulation of the rodent hippocampus at theta’s troughs induces long-term potentiation (LTP; (Huerta and Lisman, 1995; Hölscher et al., 1997; Hyman et al., 2003; McCartney et al., 2004), while stimulation during the peaks does not impact plasticity or induces depotentiation (Huerta and Lisman, 1995, 1996a; Hölscher et al., 1997; Hyman et al., 2003; McCartney et al., 2004), with consequences for memory (Siegle and Wilson, 2014; Nokia et al., 2015). Theta phase is also relevant for human memory, with information more reliably decoded at opposing phases during encoding and retrieval in intracranial hippocampal recordings (Estefan et al., 2021) and in surface EEG recordings (Kerrén et al., 2018). Perhaps the peak and trough of the saccadic ERP might likewise segregate the encoding and retrieval of visual information in the human hippocampus? Kragel et al. (2020) provided preliminary support for this idea by inferring periods of encoding and retrieval from memory-guided eye movements(Kragel et al., 2020). To date, however, direct manipulations—akin to those instrumental in uncovering causal mechanisms in rodents—have not been applied in humans to test this relationship.

Here, we assess how electrically stimulating the human hippocampus during the saccadic ERP influences memory. We built on rodent research(McCartney et al., 2004) using high-frequency electrical stimulation calibrated to the timing of the peak and trough of the saccadic ERP. We first computed saccadic ERPs from hippocampal intracranial electroencephalography (iEEG) recordings in epilepsy patients. We then used real-time eye-tracking to synchronize hippocampal stimulation with the estimated ERP peaks and troughs while participants visually explored trial-unique natural scenes with embedded targets. Subsequently, we tested their memory for each scene and associated target. Randomly timed stimulation reliably impaired memory formation compared to an unstimulated sham protocol, consistent with the bulk of hippocampal stimulation research (Lacruz et al., 2010; Jacobs et al., 2016; Goyal et al., 2018). By contrast, memory was robust to stimulation at both saccadic ERP peaks and troughs. Complementing these robust times, memory is particularly impaired by stimulation at our shortest saccade delays. Together, these results suggest that the hippocampus rapidly evolves through states that are especially sensitive or robust to stimulation following saccades..

## Materials and Methods

### Participants

Eighteen participants completed at least one memory session (40 encoding and 80 retrieval scenes) while receiving electrical stimulation. Of these participants, 14 received stimulation to their hippocampus. Other stimulation locations included the amygdala, prefrontal cortex, and parahippocampal cortex. Here, we report on those who received hippocampal stimulation, the site of our primary question All participants were implanted with Behnke Fried Macro-Micro electrodes (Ad-Tech Medical, Racine, MN, nominal diameter 1.28 mm and 5mm between contacts) to record iEEG, with locations strictly determined by clinical considerations. Participants were undergoing clinical observations at the epilepsy monitoring unit at Toronto Western Hospital to localize epileptogenic regions (Table 1). All participants voluntarily provided written informed consent, and all research was performed in accordance with protocols approved by the University Health Network Research Ethics Board.

**Table 1.** Participant Summary Table. This table groups participants according to the timing of their stimulations (see Methods): customized (Dark Grey); population average (Light Grey); partially customized white, with a * indicating marking the phase (peak or trough) with customized timing. Note that participants who received stimulation at multiple sites have a row corresponding to each site. Sessions denote the number of complete sessions (40 encoding trials and memory test), and the parenthetical recognition/target model values correspond to the number of sessions that passed inclusion criteria thresholds and were included in the recognition and target memory models, respectively. A, Anterior; HC, Hippocampus; L, Left; MTL, Mesial temporal lobe; P, Posterior; R, Right; WM, White matter.

| ID | Sex | Hand | Seizure Zone | Locatio | Cathode | Anode | Peak (ms) | Trough (ms) | Intensity (mA) | Custom (Y/N/P) | Sessions | Level of Education | Age | Onset Age |
| --- | --- | --- | --- | --- | --- | --- | --- | --- | --- | --- | --- | --- | --- | --- |
| 1 | M | R | R Temporal | RPHC | HC | WM | 149 | 236 | 3 | Y | 3(3/3) | Bachelor's Degree | 26 | 14 |
|  |  |  |  | RAHC | HC | HC | 149 | 236 | 3 | Y | 2(2/2) |  |  |  |
| 2 | F | L | R Temporal | LAHC | HC | HC | 61 | 167 | 3.5 | Y | 3(3/3) | Associate Degree | 45 | 12 |
| 12 | M | R | Not Temporal | RAHC | HC | HC | 57 | 170 | 2.5 | Y | 2(0/2) | Unknown | 49 | 0-18 |
| 13 | M | R | Not Temporal | RAHC | HC | HC | 212 | 316 | 1.5 | Y | 3(3/3) | High School | 26 | 9 |
|  |  |  |  | RPHC | HC | HC | 191 | 297 | 4 | Y | 2(2/2) |  |  |  |
| 14 | F | R | L Temporal | LPHC | HC | HC | 190 | 130 | 1.5 | Y | 3(1/0) | College | 64 | 20 |
| 22 | M | R | L Temporal | RAHC | HC | HC | 71 | 139 | 2.5 | Y | 3(2/2) | Junior High | 28 | 16 |
| 28 | M | mixed | bitemporal | LPHC | HC | WM | 92 | 145 | 2.5 | Y | 3(3/3) | High School | 28 | 13 |
| 3 | M | R | L Temporal | RAHC | HC | HC | 50 | 260 | 2.5 | N | 3(3/3) | College | 20 | 15 |
| 6 | M | R | L Temporal | RAHC | HC | HC | 50 | 260 | 2 | N | 3(1/1) | 12<sup>th</sup> grade | 27 | 7 |
| 7 | F | R | Bitemporal | RAHC | HC | HC | 50 | 260 | 4 | N | 2(1/2) | 12<sup>th</sup> grade | 58 | 14 |
| 9 | F | R | Left Temporal Not Definitive | RAHC | HC | HC | 50 | 260 | 1.5 | N | 3(1/3) | High School | 46 | 14 |
| 10 | F | R | L Insula | RAHC | HC | HC | 50 | 260 | 3 | N | 3(2/0) | High School | 20 | 6 |
|  |  |  |  | RPHC | HC | HC | 50 | 260 | 2 | N | 3(2/2) |  |  |  |
| 14 | F | R | L Temporal | LAHC | HC | HC | 60 | 130 | 2 | N | 3(2/1) | College | 64 | 20 |
| 14 | F | R | L Temporal | LPHC | HC | HC | 60 | 130* | 1.5 | P | 3(0/0) | College | 64 | 20 |
| 16 | M | R | L Temporal | RAHC | HC | HC | 78* | 130 | 1.5 | P | 3(3/3) | Associate Degree | 28 | 26 |
| 27 | M | mixed | Right temporal neocortex, not mesial temporal | RAHC | HC | HC | 78* | 130 | 3 | P | 3(3/3) | High School | 25 | 13 |
|  |  |  |  | RAHC | HC | HC | 78* | 130 | 3 | P | 3(3/3) |  |  |  |
| 28 | M | mixed | Bitemporal | RPHC | HC | WM | 60 | 131* | 2 | P | 3(2/2) | High School | 28 | 13 |

### Experimental Design and Task

#### Closed-Loop Deep Brain Stimulation System

We developed a closed-loop stimulation system which delivered electrical stimulation following online detection of eye movements (Figure 1A). We placed an EyeLink Portable Duo (SR Research Ltd., Osgoode, Canada) at the bottom of the experiment presentation laptop screen to track eye position at 1000 Hz. We used the EyeLink EDF Event Detector to pre-process eye position data and then we identified saccades using the Identification by Velocity Threshold (IVT) algorithm (>30 deg/sec) on a sample-by-sample basis as we have done before (Katz et al., 2022). The IVT has been shown to be closest to human detection of saccade events (Andersson et al., 2017). Of note, we required more than 10ms of velocity data following saccade classification to minimize the chance of triggering stimulation off eye blinks, which are often initially misclassified as saccades. Once a saccade was detected, we sent a signal to the experiment presentation software (Presentation™, Neurobehavioral Systems), which then signalled our purpose-specific Arduino to trigger an Ojemann Grass Stimulator™. This stimulator then applied customized bi-polar hippocampal electrical stimulation through implanted macroelectrodes. In tandem, iEEG data was recorded using a 256-channel Neuralynx™ Atlas Data Acquisition System (Neuralynx Inc, Bozeman, MT) at a sampling rate of 32 kHz, subsampled to 4 kHz with a high pass filter of 0.1 Hz and low pass filter of 1000 Hz. Neural and eye-tracking data were synchronized by sending TTL triggers from the experiment presentation computer to both the Neuralynx computer and stimulator using a splitter.

#### Electrode Placement

Depth electrodes were placed stereotactically using either a Leksell frame or robotically (Renishaw Neuromate). A 4-contact subgaleal electrode was used for ground and reference. This strip was placed over the parietal midline facing away from the brain. Electrode localization was performed by co-registering pre-op MRI with post-op CT using the iELVIS toolbox (Groppe et al., 2017). Only contacts that a neurosurgeon visually verified as being in the hippocampal body were used as the cathode for stimulation. Anode contact locations were restricted to the hippocampus or surrounding white matter (Table 1).

#### Stimulation Patterns and Customization

Each “electrical stimulus” consisted of a train of five 0.1ms biphasic pulses separated by 2ms. We selected this pattern since, when it is delivered at theta frequency (Larson et al., 1986) to the peak of theta following a phase-reset (McCartney et al., 2004), it optimally induces LTP (Huerta and Lisman, 1995, 1996b; Hölscher et al., 1997). During the experimental session, we applied four stimulation protocols as participants encoded information: 1) stimulation at *peak* of saccade-locked ERP; 2) stimulation at *trough* of saccade-locked ERP; 3) *sham* stimulations (no stimulation); and 4) *random*ly timed stimuli. The number of stimuli in the random condition were yoked to the participant’s saccade rate (rounded to the nearest saccade/second), ensuring that stimulation conditions only vary in the timing of the stimulation. However, they were delivered at random times throughout each second of the encoding trial using the Box-Muller transform method for sampling uniform distribution (in this case, milliseconds from each second). We applied each of the stimulation protocols across a block of 10 trials within a session to reduce carry-over effects, counterbalancing the order across participants to reduce order effects. Post hoc, we performed Wilcoxon Ranked Sum Test to confirm that stimulation types did not differ in saccade and stimulation event counts (Figure 3).

In a calibration session before the main experiment, we customized two aspects of the electrical stimuli: the saccade-locked timing and the intensity (Figure 1B). During the calibration session, participants first received task instructions and then completed an extended practice session. Mimicking the full task, they visually explored 40 scenes with subsequent testing of their recall for a subset of them (see Memory Task details below). To calibrate the timing of saccade-locked theta peaks and troughs, we obtained a saccade initiation-locked ERP for all grey matter electrodes while participants explored the 40 scenes. We then calculated the timing of statistically significant maxima (peak) and minima (trough) using polarity permutation testing (Maris and Oostenveld, 2007). We identified electrodes in the hippocampus that showed a significant peak and trough and used these times to customize the stimulation timing in the main task (*Customized Participants*). Participants who did not show a significant ERP were stimulated with the average peak and trough timing of previously tested participants (*Non-Customized*). The average times (50ms and 260ms) for non-customized participants tested early in the experiment were based on saccade locked ERP timings from our previous work (Katz et al., 2020). After a sufficient sample was collected on the current protocol (n= 9), we replaced the average times with those derived in this task’s practice session (60 and 130ms). Specifically, these times were based on the peak and trough of the average ERP across hippocampal electrodes for all participants collected to that point.

We calibrated the intensity of stimulation with a novel cortico-cortical evoked potentials (CCEPs) approach. CCEPs measure cortical connectivity (Kubota et al., 2013; Keller et al., 2014; Enatsu et al., 2015), making them a powerful index of the stimulation’s downstream effect. We, thus, inferred that the minimum current required to elicit a CCEP in any grey matter electrode was sufficient to evoke presynaptic action potentials that propagated along axons to generate the post-synaptic CCEP response. We began at 0.5mA of current for each stimulation site, increasing by steps of 0.5mA until a reliable CCEP was observed. For each current level, stimuli were delivered in a train based on the participant’s average saccade rate (rounded to the nearest saccade) during the calibration session. This stimulation was delivered for 4 seconds with 2-second breaks to match our task timing. Stimulation intensity did not exceed 8mA (Lacruz et al., 2010) as a safety precaution, and we expected less than 3mA to be sufficient (Talakoub et al., 2016; Ezzyat et al., 2017). We initially used 12 bursts of 5 pulses (Participants 1 and 2). We switched to 36 bursts to ensure reliable ERPs at lower currents for the remaining participants. The lowest current level that resulted in a significant CCEP identified with permutation testing (Maris and Oostenveld, 2007) was used for closed-loop stimulation. The average intensity across the sessions were 2.48 mA ±.80.

#### Memory Task

We developed a memory task that encourages visual exploration during learning to test the effect of saccade-locked DBS on memory formation.

##### Stimuli and Apparatus

Images of conceptually distinct outdoor and indoor scenes were used as stimuli in the memory task. They were collected using Google image search and a database of scenes (Zhou et al., 2018) standardized to 1400x1000 pixels. Four copies of a target image – either a cartoon bunny or coyote – were positioned within each of the scenes according to an updated version of the universal image quality index (UIQI; Wang and Bovik 2002) called the Structural Similarity Index (SSIM) (Wang et al., 2004). The algorithm compares the target’s similarity to each possible placement within the scene to find locations where it would be camouflaged, and UIQI has been shown to increase visual search times in camouflage (Lin et al., 2014).

The experiment was presented on a 15-inch laptop using Presentation software (NeuroBehavioral Systems, Albany, CA, USA). Participants were seated in their hospital beds, and the laptop was placed in front of them at a comfortable viewing distance. An infrared video-based eye tracker (EyeLink Duo; SR Research Ltd., Osgoode, Canada), connected to the laptop with ethernet, was used to monitor and record their eye movements throughout the task. All participants first underwent a 13-point eye-tracker calibration and 9-point validation test before starting each day of testing.

##### Procedure

Each memory task session consisted of an encoding and a retrieval phase. In the study phase, participants were presented with 40 scene images embedded with four copies of one of two possible targets (Figure 1C). Participants were instructed to visually search through the image to find as many of the targets as possible. To increase the depth of their learning, they were instructed that if they found all four targets to keep exploring the scene and elaborate on what the cartoon character would do in the scene. Each image was presented for four seconds, after which participants reported how many targets they found. When it did not interfere with the quality of eye-tracking data, participants used a keyboard to make responses (27/59 sessions); otherwise, they responded verbally, and the experimenter pressed the corresponding key. Each response was followed by a 1-second fixation cross, followed by the next image. To help participants build associations between scene images and targets, each target was repeated for alternating blocks of 5 trials. Across participants, scene images were consistently paired with the same targets and were presented in the same order within a session. Stimulation type was counterbalanced across participants, such that a particular stimulation type was similarly likely to be applied in each quartile of the encoding session and to each studied scene image.

Immediately following the encoding phase, participants were presented with instructions and the retrieval phase task was administered. On each of 80 retrieval trials, participants were first presented with either a scene from the preceding encoding phase or a new, unstudied scene. Target images were not embedded in these scenes. Participants were first asked to indicate whether they recognized the scene from the studied set or if they thought it was new. They then rated their confidence in their recognition judgement on a scale from not sure (1) to very sure (4). Lastly, if they recognized the scene, they were asked to recall which target was embedded within it. Participants were given unlimitted time to answer each question. This task was broken into blocks of three self-contained sessions. If time allowed, different stimulation locations (e.g., anterior or posterior hippocampus) were tested in a second block.

**Figure 1.**
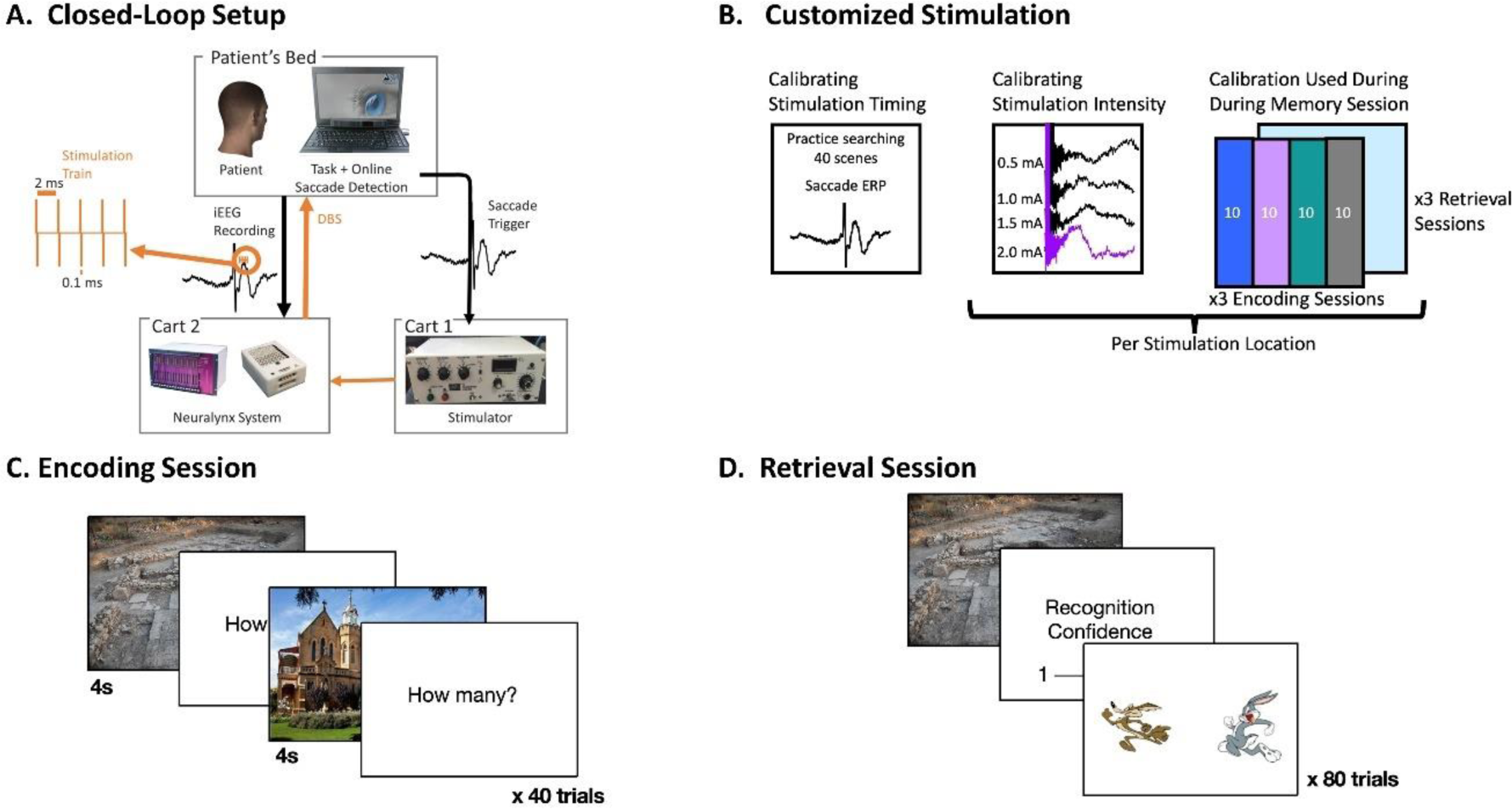
Experiment Design: **A)** Block diagram of the closed-loop stimulation system. Stimulation bursts (orange; 5 pulses; 500 Hz; 0.1ms pulse width) were applied following the online detection saccades to coincide with the resulting ERP peak or trough. **B)** Stimulation calibration protocol. Participants practiced 40 encoding trials during which a saccade-locked ERP was extracted for customized stimulation timing. Then, during the calibration session, stimulation bursts were applied according to the participant’s saccade rate from the practice session to determine the minimum intensity needed to elicit a significant response in non-stimulated grey matter electrodes (2.0mA in the example). The rightmost schematic illustrates an example counterbalancing scheme for stimulation types (different color panels) during a session. **C)** Example encoding session trial. **D)** Example retrieval trial, including scene recognition judgement, confidence rating, and associative memory probe for the target.

### Data and Statistical Analysis

#### Behavioural Data Analysis

We used a generalized linear mixed-effect model (lme4 package; version 1.1.26, (Bates et al., 2014)) in R programming language (version 1.4.1106; to predict subsequent scene recognition (hit rate) and associative target memory accuracy on a trial-by-trial basis according to the trial’s stimulation type. Such models are recommended to account for non-independence of trials within participants and to account for random effects when dealing with intracranial brain stimulation and declarative memory (Suthana et al., 2018). We included a random slope for stimulation type and random intercept grouped by participant. We used *a priori* data exclusion criteria to restrict analyses to sessions with reliable memory performance. For the scene recognition model, sessions were excluded if they had corrected recognition (hit rate – false alarm rate) scores lower than 0.25; one subject and 17 sessions were excluded based on this criterion. We also excluded recognition judgements with reaction times greater than 10 seconds. For the associative memory model, sessions were further excluded if less than 15 old scenes were recognized because participants only made associative target judgements on the scenes that they recognized; 16 additional sessions were removed based on this criterion. We further excluded target memory responses longer than 5 seconds; this cut-off was shorter than for recognition judgments because, unlike recognition judgments, the relevant scene was not visible to guide retrieval during this decision.

To identify potential moderators of stimulation type on memory, we separately included the interaction of stimulation type (peak, trough, random, sham) with various covariates in distinct models. These covariates included: location within the hippocampus (anterior, posterior); laterality (left, right); the number of stimulation events per trial; timing customization; stimulating in the seizure onset zone; and stimulation intensity. Stimulation intensity was centered across participants, and number of stimulations was centered within participant. Most of these covariates did not influence the impact of stimulation type. There was a marginal interaction between seizure onset zone and stimulation type, where the impact of trough stimulation was marginally less detrimental to scene memory formation when applied in the seizure onset zone than outside of it (β = 0.74, SE = 0.43, *t* = 1.72, *p* = 0.09). However, since only stimulation customization significantly moderated the impact of stimulation type (see Results), the final model was:

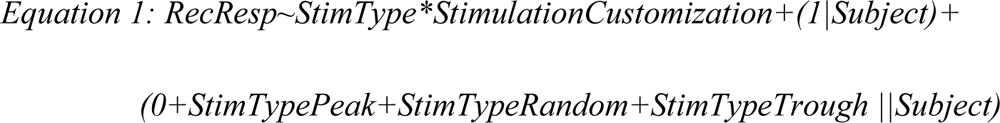

The emmeans package (version 1.5.5.1) was used to perform follow-up simple effect comparisons when motivated by an interaction between fixed effects. Lastly, since stimulation customization proved to be an important factor, we also ran a model predicting sham memory performance based on customization group (with random intercepts per participant) to determine if memory differed between groups when no stimulation was applied.

Lastly, we used a mixed linear model to ask how the raw timing of stimulation (regardless of customization and peak/trough label) influenced memory. This analysis was restricted to peak and trough protocols because they were the only conditions to use consistent saccade-locked timing within a trial. Subsequent recognition accuracy (hit rate) for each stimulation session was corrected by subtracting each participant’s hit rate from that session’s sham condition. This corrected hit rate was then linearly predicted by the delay between saccade initiation and stimulation. The model was restricted to stimulation conditions applied <150 ms following a saccade, which resulted in one or two corrected hit rates per session, depending on the timing of peak and trough stimulation (see Table 1). Note that, since this analysis assessed relationships between continuous variables, outlying points—identified as having Cook’s Distances above 4/n—were removed.

#### iEEG Data Analysis

All electrophysiological data was pre-processed by downsampling each trial to 1 kHz, bandpass filtering between 0.5 and 200 Hz using a second-order Butterworth filter, and notch filtering at 60 Hz to remove line noise. Notch filters were also incorporated at harmonics of 60 Hz up to 200 Hz. All further analysis was performed using custom-written scripts in MATLAB (The Mathworks Inc, Natick, MA, USA).

To obtain ERPs for saccade or stimulation onset events, iEEG data from all trials and all relevant electrodes were aligned to the initial eye movement or electrical stimulation-onset and trimmed to epochs containing 1.2 seconds of data before and after each event. The ERP for each electrode was then obtained by taking the mean of these saccade-onset epochs across all trials from all experimental blocks.

To determine the statistical significance of each ERP, a surrogate distribution of ERP maxima and minima was obtained for each electrode/ERP using non-parametric permutation testing with randomized polarity inversions (3000 permutations). ERP values were deemed significant if they fell above the 97.5^th^ percentile of the distribution of maxima or below the 2.5^th^ percentile of the distribution of minima.

## Results

During 59 sessions, we stimulated in 19 hippocampal locations (Figure 2A) across 14 participants. We leveraged the robust hippocampal ERPs elicited by eye movements (Katz et al., 2020) to stimulate the hippocampus during different post-saccade electrophysiological states. For contacts that showed statistically significant saccadic ERP peaks or troughs during the practice session, we customized stimulation timing (fully customized: 24/59 sessions; peak only customized: 9/59 sessions; trough only customized: 6/59; Figure 2B). For sessions where the corresponding peak or trough did not reach statistical significance, the *Non-Customized* average timing (Figure 2C) was used (fully non-customized: 20/59 sessions). To verify the timing of our customization, we visualized when stimulation would have been delivered with respect to the saccadic ERP from all sham trials – the trials with no stimulation artifact. As visualized in an example participant in Figure 2D, the peak and trough times calibrated during the practice session remained aligned to the peak and trough of the saccadic ERP during experimental sessions.

**Figure 2.**
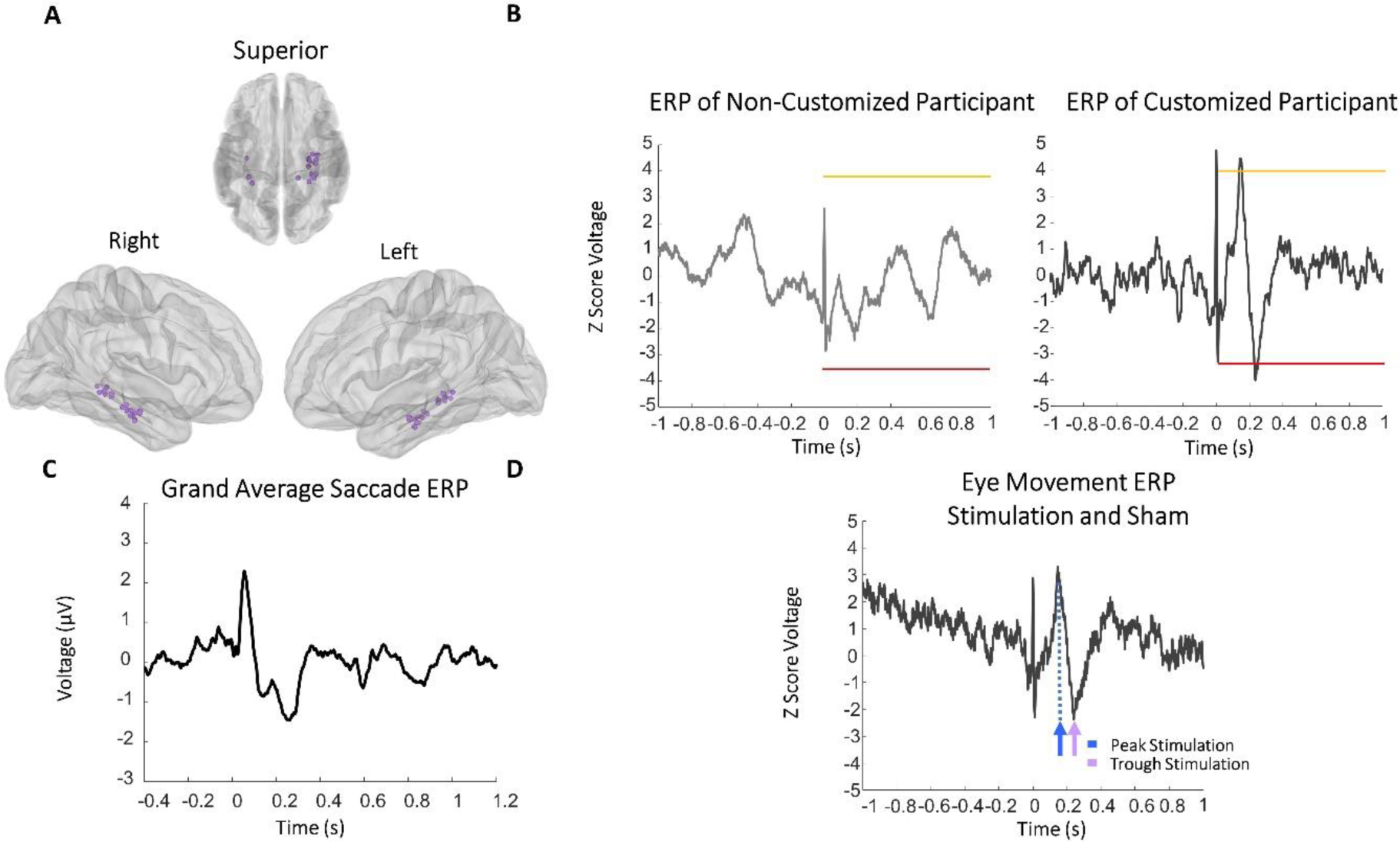
Extracting timing of stimulation. **A)** Demonstrates the location of the cathode for electrical stimulation. Each colour is a different subject for that hippocampal location*. **B)** Example saccade-locked ERP from Non-Customized and Customized timing participants recorded during the practice session. The yellow and red bars mark statistical thresholds. **C)** When no significant ERP was elicited in a *Non-Customized* participant, the stimulation were based on group-average ERPs (peak=50ms, trough=260ms for first 9 participants; peak=60ms, trough=130ms for later participants, shown in figure). **D)** Verification of customized timing. Example customized participant saccade-locked ERP from sham trails. Arrows designate peak (blue) and trough (purple) stimulation times calibrated during the practice session, demonstrating that the customized timing remained accurate throughout the stimulation sessions. ***\*****Participant 2’s stimulation location is omitted because it was corrupted on the research files which enable visualization in this capacity*

### Eye movements and stimulation

We confirmed that stimulation protocols did not differ in their rate of stimulation events. Specifically, the number of stimulation events applied during the sessions included in recognition analyses did not significantly differ between peak, trough, and random protocols (|*z*| < 1.53, all *p* > 0.127, uncorrected; Figure 3a). This consistency was because the saccade rates also did not differ across all four protocols (|*z*| < 1.65, all *p* > 0.100, uncorrected; Figure 3b). Participants made an average of 3.65 ±0.59 saccades/second across all the sessions that were included in the recognition memory model.

**Figure 3.**
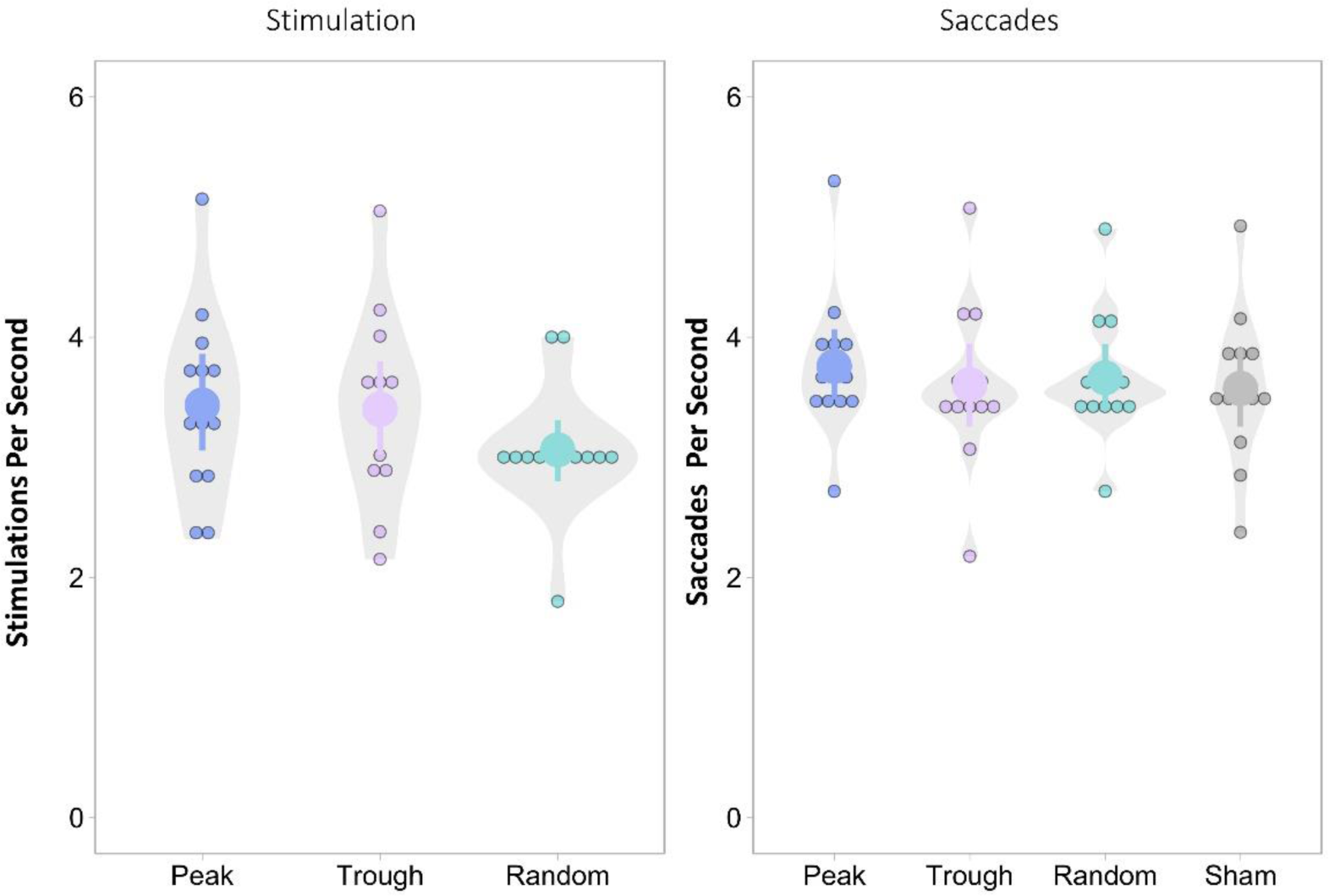
Stimulation and Saccade Event Distributions Based on Stimulation Type: **A)** Number of stimulation events administered per session did not significantly differ across stimulation types. **B)** Number of saccades produced also did not differ between different stimulation types. Small dots indicate average rate per participant; large dots indicate the group average and surrounding lines reflect the bootstrapped 95% confidence intervals.

### Overall Memory Performance

Overall, 4.21 ± 2.01 sessions were performed by each participant. An average of 62.5% (SE=8%) of old scenes were recognized (hits), and 22% (SE=5%) of new scenes were falsely endorsed as being studied (false alarms), leading to an average corrected recognition rate of 0.404, which was well above chance (*t*(13) = 7.4, *p* < 0.5*10^-5^). Associative memory was not above chance across all participants 54.0% (t(13) =1.5976, *p* = 0.134). However, when the analysis was restricted to those with a sufficient number of associative judgements, performance was above chance (54.87%; t(13) = 2.52, *p* = 0.026). We also confirmed that fully customized and non-customized participants did not significantly differ in their memory performance for sham (baseline) trials (Recognition: β = -0.042, SE = 0.085, *t* = -0.492, *p* = 0.627; Associative memory: β = 0.056, SE = 0.08, *t* = 0.70, *p* = 0.50) or in their customized stimulus intensity (β = 0.33, SE = 0.42, *t* = 0.79, *p* = 0.44). Likewise, the intensity of stimulation was not associated with sham memory performance (Recognition: β = 0.0525, SE = 0.0428, *t* = 1.227, *p* = 0.225; Target memory: β = 0.0248, SE = 0.0476 *t* = 0.522, *p* = 0.603). Thus, the impact of these stimulation parameters on memory is more likely to reflect their direct impact on memory rather than any pre-existing differences that influenced their customization.

### Effect of stimulation types on memory across the group

Across the 42 sessions (13 participants) included in the recognition memory model, stimulation appeared to hinder memory performance (Figure 4). Participants were less likely to later recognize scenes that were studied during peak (β=-0.46, SE=0.17, z= -2.8, p=0.006) and random stimulation (β=-0.70, SE=0.17, z =-4.2, p=2.8*10-5) as compared to sham. Trough stimulation similarly trended towards decrements (β=-0.42, SE=0.23, z= -1.9, p=0.063). All stimulation types resulted in significantly worse memory than sham when all sessions were included in the model (See Supplemental Data 4B). Consistent with prior research (Lacruz et al., 2010; Jacobs et al., 2016; Goyal et al., 2018), these results suggest that applying electrical stimulation to the hippocampus disrupts memory formation, regardless of its timing. By contrast, stimulation did not influence the likelihood of participants later being able to recall the associated cartoon character, with no difference when comparing sham stimulation to peak (β=-0.13, SE=0.16,z=-.81, p=0.41), trough (β=-0.034, SE=0.16, z=-.20, p=0.84), or to random (β=-0.002, SE=0.17, z= .01, p=0.99) (Figure 4). However, this lack of effect could reflect near floor performance and low statistical power, as only a small number of trials and sessions were included in this analysis. Accordingly, we focussed on recognition memory for the remainder of the analyses.

**Figure 4.**
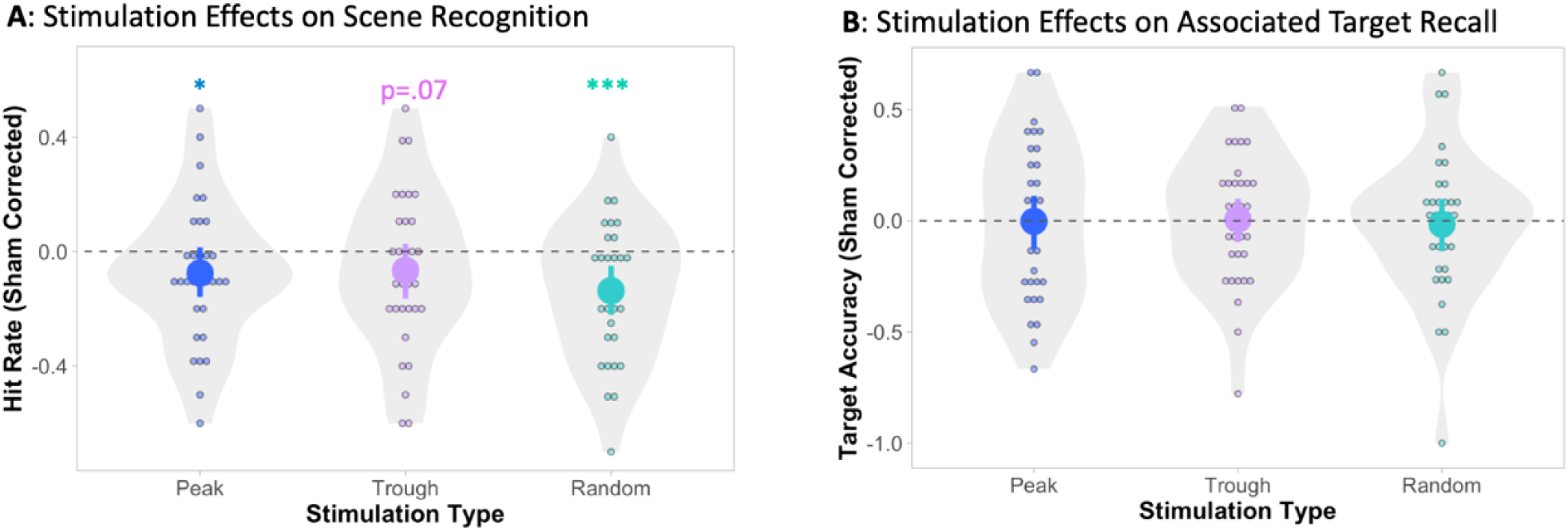
Group memory performance: Average hit rate (A) and target memory accuracy (B) for each stimulation condition when sham is used as a baseline. Small dots indicate average rates per session; large dots indicate the group average and surrounding lines reflect the bootstrapped 95% confidence intervals. Note that data is averaged within the session and sham corrected for ease of visualization, but reported statistics reflect interactions in the associated mixed model, which was estimated across trials. * p < 0.05; *** p < 0.0001

### The moderating effect of timing customization

We next determined whether each stimulation protocol’s effect on recognition memory was moderated by the specificity of its timing, i.e., whether stimulation was delivered at Customized peak and trough times or simply the Non-Customized average times. Specifically, we included customization (fully non-customized vs. fully customized) and stimulation type and their interaction in one model predicting subsequent recognition memory (Figure 5). Much like in the full sample, we found that stimulation compromised memory formation during Non-Customized sessions: Non-Customized participants were less likely to later recognize scenes presented with peak (β = -0.90, SE = 0.28, z = -3.2, p = 0.0012) or random stimulation as compared to sham stimulation (β = -0.85, SE = 0.28, z = -3.07, p = 0.0022), with a trend for the trough comparison (β = -0.69, SE = 0.41, z = -1.71, p = 0.087). However, there was a significant interaction between customization and stimulation type. Specifically, the impact of peak stimulation depended on whether its timing was customized (β = 0.96, SE = 0.38, *z* = 2.55, *p* = 0.011). This interaction was driven by the timing of the stimulation mattering for the Customized group’s memory; stimulation that was precisely timed to the peak yielded stronger memories than random stimulation (β=0.54, SE=0.25, *z* =2.22, *p* = 0.026), and precisely timed trough stimulation trended toward showing similar benefits (β = 0.67, SE = 0.39, *z*= 1.71, *p* = 0.087). Like Non-Customized participants, Customized participants were significantly less likely to recognize images encoded under random as compared to sham stimulation (β = -0.48, SE = 0.24, *z* = -1.99, *p* = 0.047). We also ran this model including Partially Customized participants, labeling each peak and trough stimulation according to its respective customization. See Figure 5 Supplement for a similar pattern of results. Thus, applying stimulation to the hippocampus tends to disrupt memory formation unless it is precisely aligned to physiologically meaningful states following a saccade.

**Figure 5.**
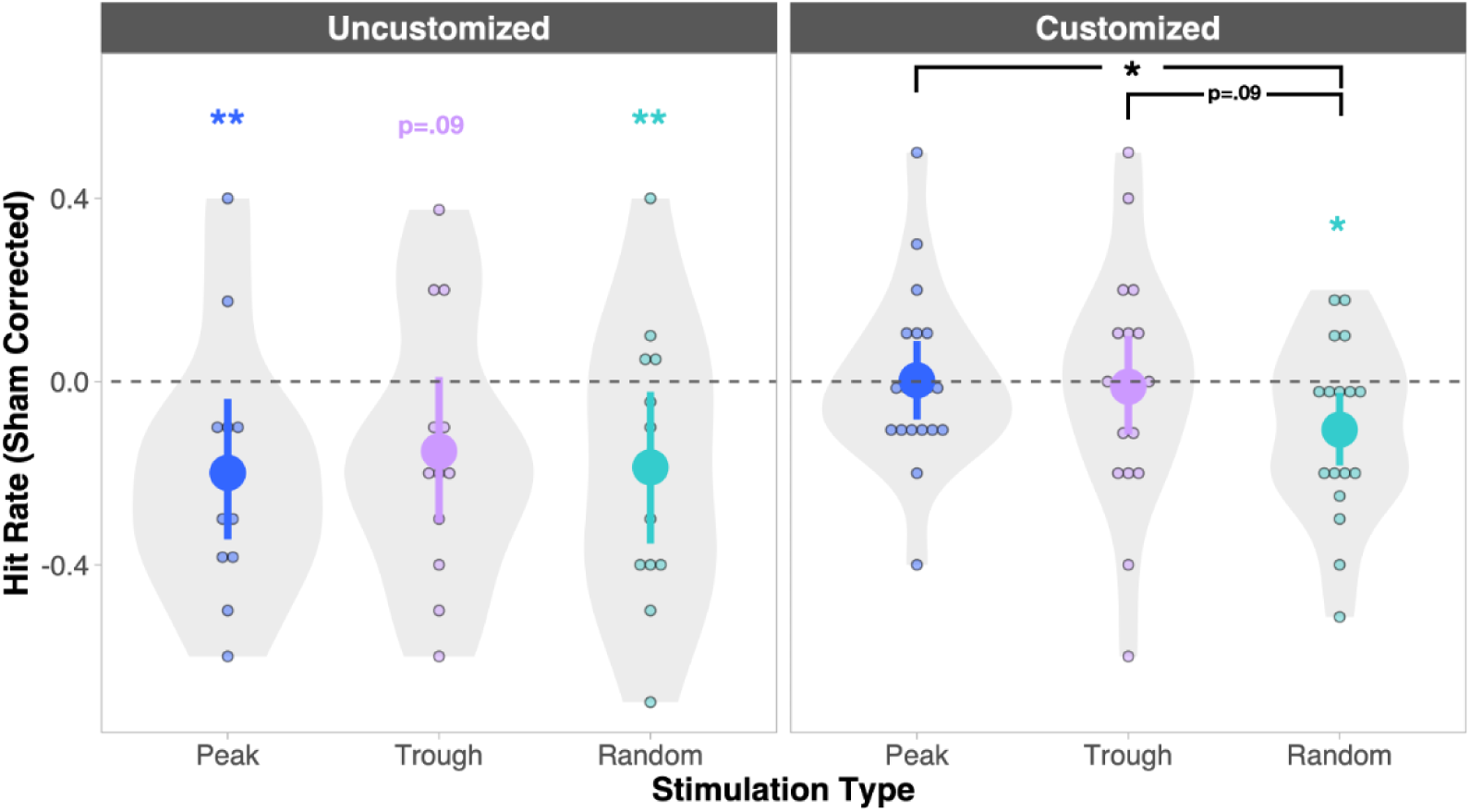
The moderating effects of customized stimulation timing on recognition memory: Average hit rate for *Non-Customized* (left) and Customized (right) sessions. Sham is used as a baseline for all stimulation types. Small dots indicate average rates per session; large dots indicate the group average and surrounding lines reflect the bootstrapped 95% confidence intervals. Note that data is averaged within sessions and sham corrected for ease of visualization, but reported statistics reflect interactions in the associated mixed model and follow-up contrasts performed with emmeans, which were estimated across trials. Colored significance markers denote differences from sham (baseline); markers contained in black bars mark differences between active stimulation conditions. Note that this analysis was motivated by the significant interaction between customization and stimulation type (peak vs. sham). * p < 0.05; ** p < 0.005

### The effect of raw stimulation timing

Lastly, we explored if there were particularly disruptive times to stimulate the hippocampus with respect to eye movement initiation. We recently characterized how individual medial temporal lobe neurons change their firing rates around saccades (Katz et al., 2022). We found that putative excitatory units decreased their firing rates during the perisaccadic interval, while putative inhibitory units increased their firing rates. The net effect of this pattern likely reflects a period of inhibition starting shortly before the initiation of the saccade and lasting for around 100 ms. Accordingly, we asked if stimulating during this early, inhibitory phase would especially impair memory. Leveraging the natural variability in stimulation timing across people, we analyzed subsequent memory performance as a function of stimulation timing, focussing on times that were under 150 ms from the saccade onset. We found a roughly linear relationship between memory and stimulation time, with performance steadily increasing with the delay from saccade initiation (β = 0.002, SE = 0.0008, *t*(34.8) = 2.52, *p* = 0.02; Figure 6). Notably, this linear relationship did not extend to later delays when the same model extended to include all peak and trough sessions (β = 0.0003, SE = 0.0003, *t*(58.4) = 0.85, *p* = 0.40). The relation was also unlikely to be driven by customization as there was a mix of customized and non-customized sessions at early and late delays. Instead, it was likely driven by images being particularly hard to remember when electrical stimuli fell within the inhibitory period that follows saccades.

**Figure 6.**
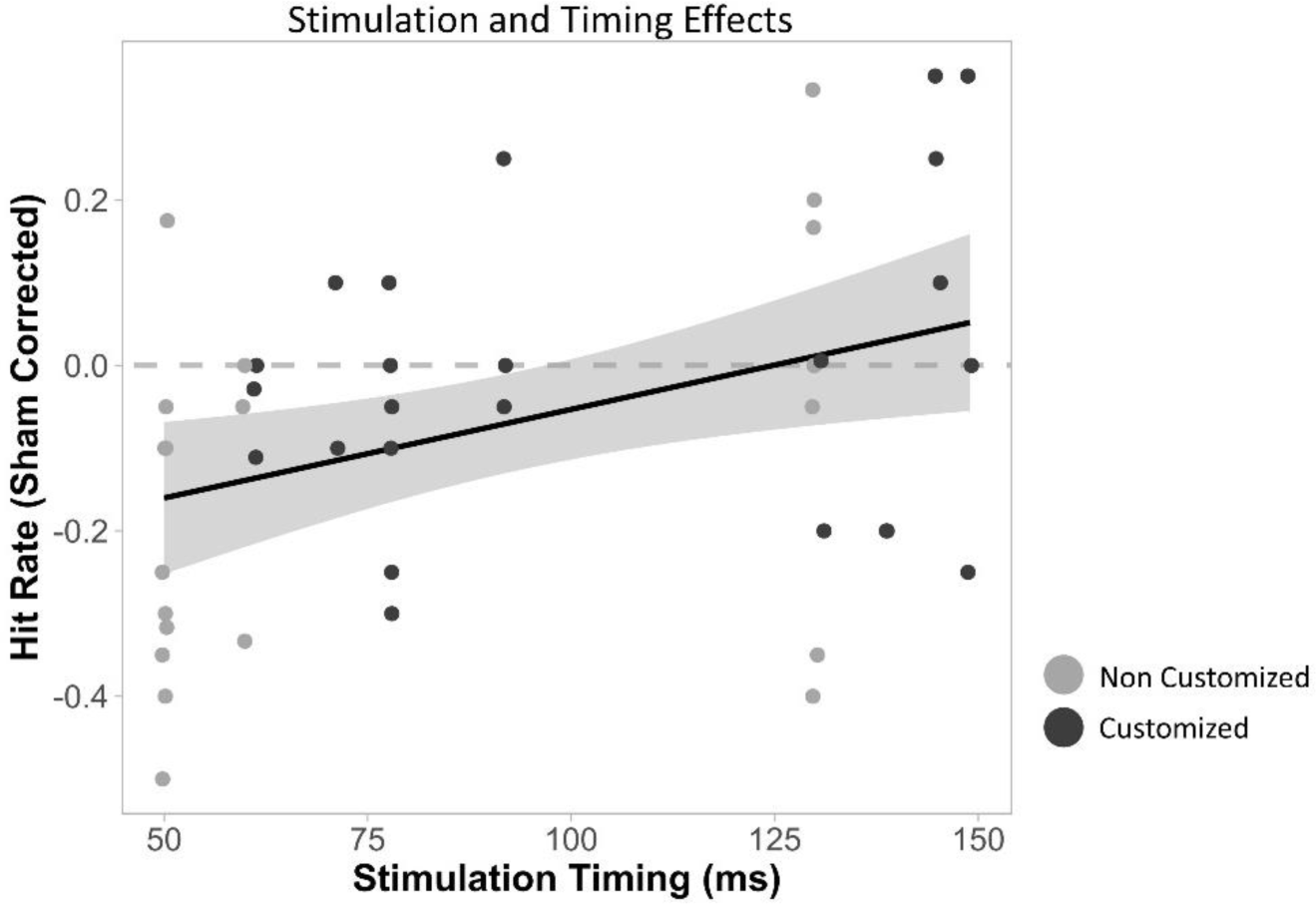
Subsequent recognition depends on raw stimulation timing. Y-axis depicts Sham corrected subsequent hit rates for sessions (dots) as a function of the stimulation timing (relative to saccade onset) used during encoding. Note the mix of customized(dark) and non customized (light) sessions at shorter and longer stimulation delays.

## Discussion

Why does visual exploration improve memory formation (Meister and Buffalo, 2016)? Here, we explore the possibility that eye movements create physiological states relevant for memory formation, analogous to those created by resetting hippocampal theta phase (Hasselmo et al., 2002; McCartney et al., 2004). We timed hippocampal electrical stimulation to target the peak and trough of people’s saccade-related ERPs while they formed memories. Stimulation generally impaired subsequent recognition memory without altering the number of eye movements produced. However, memory recovered to baseline levels when peak and trough were precisely targeted, suggesting that the generating brain states are similarly robust to perturbation. By contrast, stimulating within 100 ms of saccade initiation—when the MTL is usually inhibited (Katz et al., 2022)—particularly disrupted later recognition. Thus, the hippocampus may rapidly cycle through states that are robust and vulnerable to perturbations following each eye movement.

The recognition memory impairments observed in most stimulation conditions align with and extend the literature. Prior work has also reports memory formation impairments when stimulating the hippocampus (Halgren and Wilson, 1985; Halgren et al., 1985; Coleshill et al., 2004; Jacobs et al., 2016; Merkow et al., 2016; Goyal et al., 2018; Kucewicz et al., 2018b, 2018a). However, these studies stimulate the hippocampus continuously for seconds, whereas we used 10 ms bursts, leaving it unmodulated for most of the encoding period. The marked impairments caused by brief bursts are even more striking when considering their timing.

Specifically, since participants saccade around 4 times per second, peak and trough protocols approximated theta-burst stimulation (TBS). TBS can improve human memory formation (Miller et al., 2015; Titiz et al., 2017; Inman et al., 2018; Mankin et al., 2021), perhaps by eliciting endogenous plasticity mechanisms (Larson et al., 1986). Differences in the tissue targeted may explain why our stimuli did not also enhance memory. Locations connected to the hippocampus—not hippocampal grey matter—were stimulated in studies showing enhancements (Miller et al., 2015; Titiz et al., 2017; Inman et al., 2018; Mankin et al., 2021). Direct stimulation of the hippocampus in protocols like ours may produce “informational lesions” because hippocampal neurons are driven by artificial (and arbitrary) stimulation rather than their (informative) inputs (Lowet et al., 2022) (Hampson et al., 2018). Thus, we add to mounting evidence that hippocampal stimulation impairs memory formation (Lacruz et al., 2010; Jacobs et al., 2016; Goyal et al., 2018) by showing that even brief bursts are sufficient to disrupt the fidelity of encoding.

In contrast to these near ubiquitous memory impairments, the hippocampus was robust to direct stimulation at the peak and trough of the saccadic ERP. But these robust states seem to be short-lived. Specifically, they were only observed when precisely calibrating to participants’ own peak and trough, and not when using the group-averaged times for Non-Customized participants. While informative, the observed group interaction was not by design. In fact, we included Non-Customized participants to maximize data collection, reasoning that group timing would approximate their (hard to measure) peak and trough. But the impairments suggest that group timing missed the short-lived mark. Alternatively, since only Customized sessions showed more reliable saccadic ERPs, they may be the only ones with robust enough states at the peaks and troughs to withstand the impact of stimulation. These possibilities could be directly arbitrated by varying the precision of stimulation timing in sessions with strong saccadic ERPs.

Why are the peaks and troughs of saccadic ERPs robust to stimulation? We draw inspiration from the prominent SPEAR model (Hasselmo et al., 2002; Hasselmo and Stern, 2014), reasoning that the phases of hippocampal saccadic ERPs—suggested to reflect a theta reset (Hoffman et al., 2013; Jutras et al., 2013; Katz et al., 2020)—may share properties with the phases of better-studied theta rhythms. The SPEAR model proposes that encoding and retrieval tend to occur during the trough and peak of hippocampal theta, respectively. Accordingly, we predicted that stimulating saccadic ERP peaks would leave memory formation relatively unchanged if these peaks likewise corresponded to periods of retrieval. Our prediction was confirmed. Conversely, we predicted that trough stimulation would influence encoding: enhancing memory if the stimulus timing fostered meaningful plasticity or impairing it if stimulation created “informational lesions.” This prediction was not supported. Perhaps the divergent possible effects—enhanced plasticity and informational lesions—canceled out. Yet, a more parsimonious explanation would be that, unlike the phases of rodent hippocampal theta, the peaks and troughs observed during human visual exploration are not differently tuned to encoding and retrieval. Before embracing this conclusion, though, we mechanistically examine the assumptions behind our initial reasoning.

First, our initial predictions naively assumed that subsequent memory *only* depends on encoding operations. A more nuanced proposal argues that strong memories derive from rapidly switching between encoding and retrieval during visual exploration, as people generate and test predictions about the scene (Kragel and Voss, 2022). More abstractly, participants might bolster their learning of commonplace environments by retrieving related experiences and using this knowledge to scaffold their memory (Gilboa and Marlatte, 2017; Bellana et al., 2021). If participants relied on both encoding and retrieval to form robust memories, stimulating during retrieval (peak) and encoding (trough) phases could yield similar subsequent memory outcomes in our task.

Alternative insights come from the mechanisms motivating the SPEAR model. Starting with circuit configurations, inputs from area CA3 vs. entorhinal cortex predominately drive CA1 neurons at the peak and trough of theta, respectively. If the peak and trough of the saccadic ERPs and theta share similar circuit configurations, perhaps the dominance of individual pathways at these moments create particularly stable circuits compared to the transitions between motifs. We recently identified such a parallel, inspired by the corollary discharge-like properties of hippocampal saccadic responses (Katz et al., 2022). Saccadic corollary discharges involve circuits that run from the superior colliculus to the thalamus, to the inhibited cortical region (Sommer and Wurtz, 2002; Berman et al., 2017). Following this pattern, the thalamic nucleus reunions could likewise relay saccade planning signals to neurogliaform (NGF) inhibitory interneurons in area CA1 (Dolleman-Van Der Weel et al., 1997; Krout et al., 2001; Morales et al., 2007; Eleore et al., 2011; Scheel et al., 2020). Crucially, NGF interneurons selectively disengage entorhinal input to area CA1, allowing CA3 input to dominate (Sakalar et al., 2022). This mirrors how medial septum neurons—which set the pace of hippocampal theta—drive interneurons to shift CA1 toward CA3 and away from entorhinal input (Hasselmo et al., 2002; Leão et al., 2012). Thus, a thalamic relay engaged during saccade initiation could generate circuit motifs that parallel those produced across theta phases; motifs that may be particularly robust to perturbation.

While shifts in synaptic plasticity across theta-phases are harder to map onto the saccadic ERP, new discoveries warrant their discussion. The proposal that Schaffer collateral LTP is best induced at theta’s troughs was based on studies that applied multiple bursts of TBS, beginning at either the trough or peak of ongoing rhythms. More recently, however, Law and Leung (Law and Stan Leung, 2018) induced LTP using a single high-frequency burst (5 pulses, like in our protocol) at many phases of mouse hippocampal theta. Rather than one optimal phase, they identified two: one at the rising phase of theta and one at the falling phase. Notably, the electrical burst did not elicit LTP at either the peak or trough of theta. This raises the intriguing possibility that saccadic peaks and troughs were so robust to stimulation because they were periods of *low plasticity,* leaving the patterns of neural activity encoded during non-stimulation times free of noise. However, much work is needed to determine how plasticity changes across the phases of the saccadic ERP.

More broadly, understanding how DBS timing influences memory outcomes has clinical implications (Kocabicak et al., 2015). Indeed, closed-loop stimulation can boost memory by stimulating the brain at the right time for memory (Ezzyat et al., 2018; Hampson et al., 2018). We add to this work by targeting two theoretically motivated points on the saccadic ERP. A more data-driven exploration of more points across the full ERP might uncover key times for stimulation that likewise benefit memory. Alternatively, using saccadic-timing to deliver stimulation to areas connected to the hippocampus formation (Miller et al., 2015; Titiz et al., 2017; Inman et al., 2018; Mankin et al., 2021)—rather than hippocampal grey matter—could benefit memory by amplifying modulatory inputs when the hippocampus is primed to receive them. Indeed, local theta phase modulates the amplitude of human hippocampal potentials evoked by stimulating connect regions (Lurie et al., 2022), suggesting that hippocampal responsiveness to input varies across hundreds of milliseconds. Thus, the saccadic stimulation protocol presented here provides another avenue in the pursuit of alleviating memory disorders.

## Conclusion

In summary, we charted how direct hippocampal stimulation influences human memory formation at critical moments following a saccade. Our results suggest that memory is particularly vulnerable to stimulation immediately following saccade initiation, potentially reflecting the consequence of exciting hippocampal neurons during what is usually an inhibited period (Katz et al., 2022). Critically, this vulnerable period was followed by two states in which memory was particularly robust to stimulation—precisely at the peak and trough of participants’ personal saccadic ERPs. These discoveries, along with our approach, have important implications for understanding the timing of human memory. Specifically, hippocampal temporal dynamics have largely been conceived through the lens of rodent physiology, which shows well-defined, continuous theta rhythms (Buzsaki et al., 1985; Buzsáki, 2002). By contrast, theta is more ephemeral in the primate hippocampus (Green and Arduini, 1954; Stewart and Fox, 1991; Bush et al., 2017; M. Aghajan et al., 2017), which instead shows pronounced saccadic responses (Sobotka and Ringo, 1997; Hoffman et al., 2013; Jutras et al., 2013; Andrillon et al., 2015; Katz et al., 2020; Mao et al., 2021). Accordingly, the saccadic responses that steadily punctuate hippocampal physiology may be a more meaningful timekeeper for primates. By causally assessing how human memory is perturbed—or rescued—across this saccadic response, we hope to inspire others to mechanistically explore how visual exploration coordinates human memory.

## Supplemental Data

**Figure 1 Supplement Related to Figure 4.**
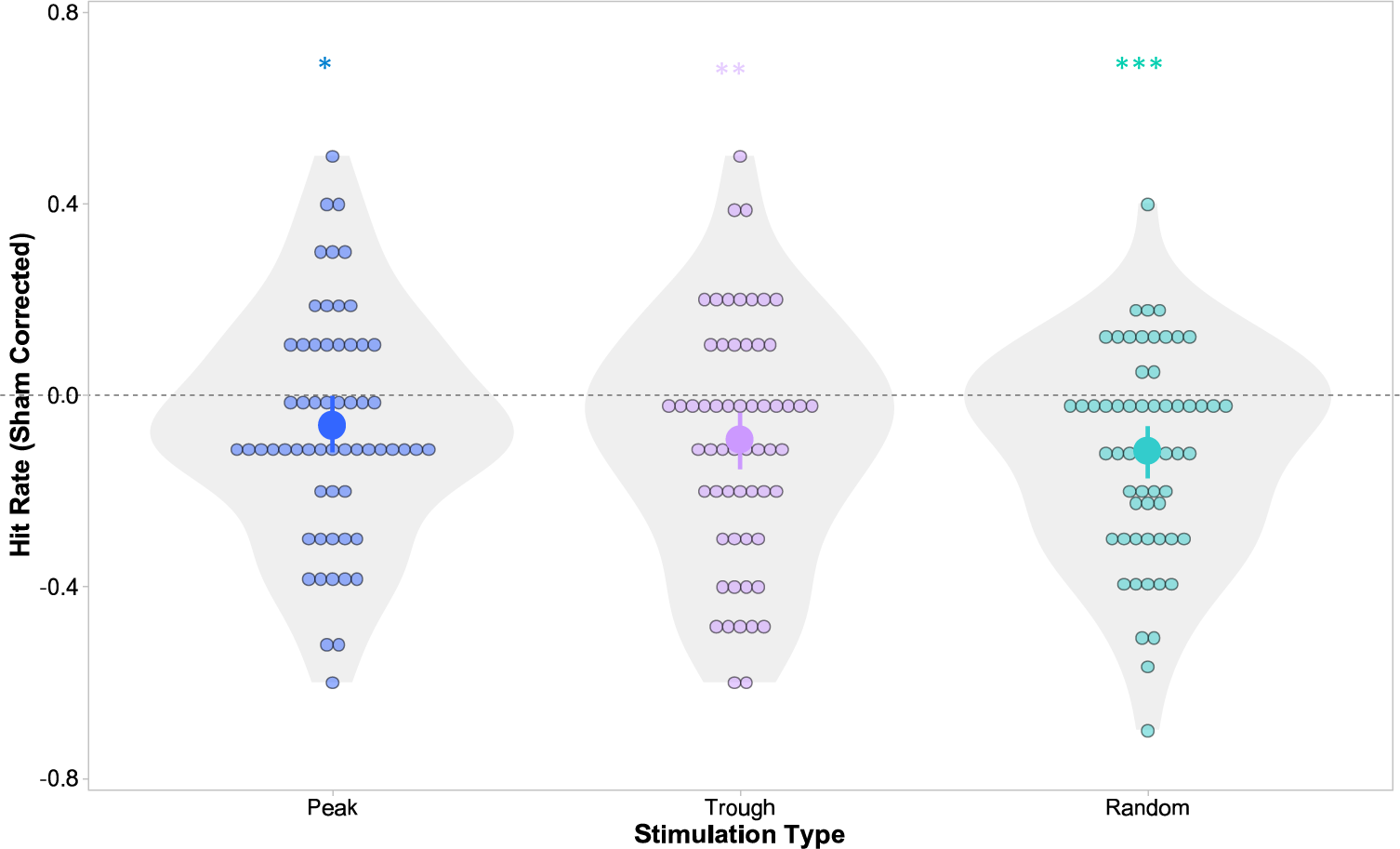
Group memory performance: Average hit rate (A) for each stimulation condition when sham is used as a baseline. Small dots indicate average rates per session; large dots indicate the group average and surrounding lines reflect the bootstrapped 95% confidence intervals. Note that data is averaged within the session and sham corrected for ease of visualization, but reported statistics reflect interactions in the associated mixed model, which was estimated across trials. Participants were less likely to later recognize scenes that were studied during peak (β=-0.35, SE=0.16, z= -2.2, p=0.027), trough (β=-0.55, SE=0.17, z= -3.16, p=0.0016), and random stimulation (β=-0.62 SE=0.14, z=-4.46, p=8.3*10-6) as compared to sham. * p < 0.05; ** p < 0.01; *** p < 0.0001.

**Figure 2 Supplement Related to Figure 5.**
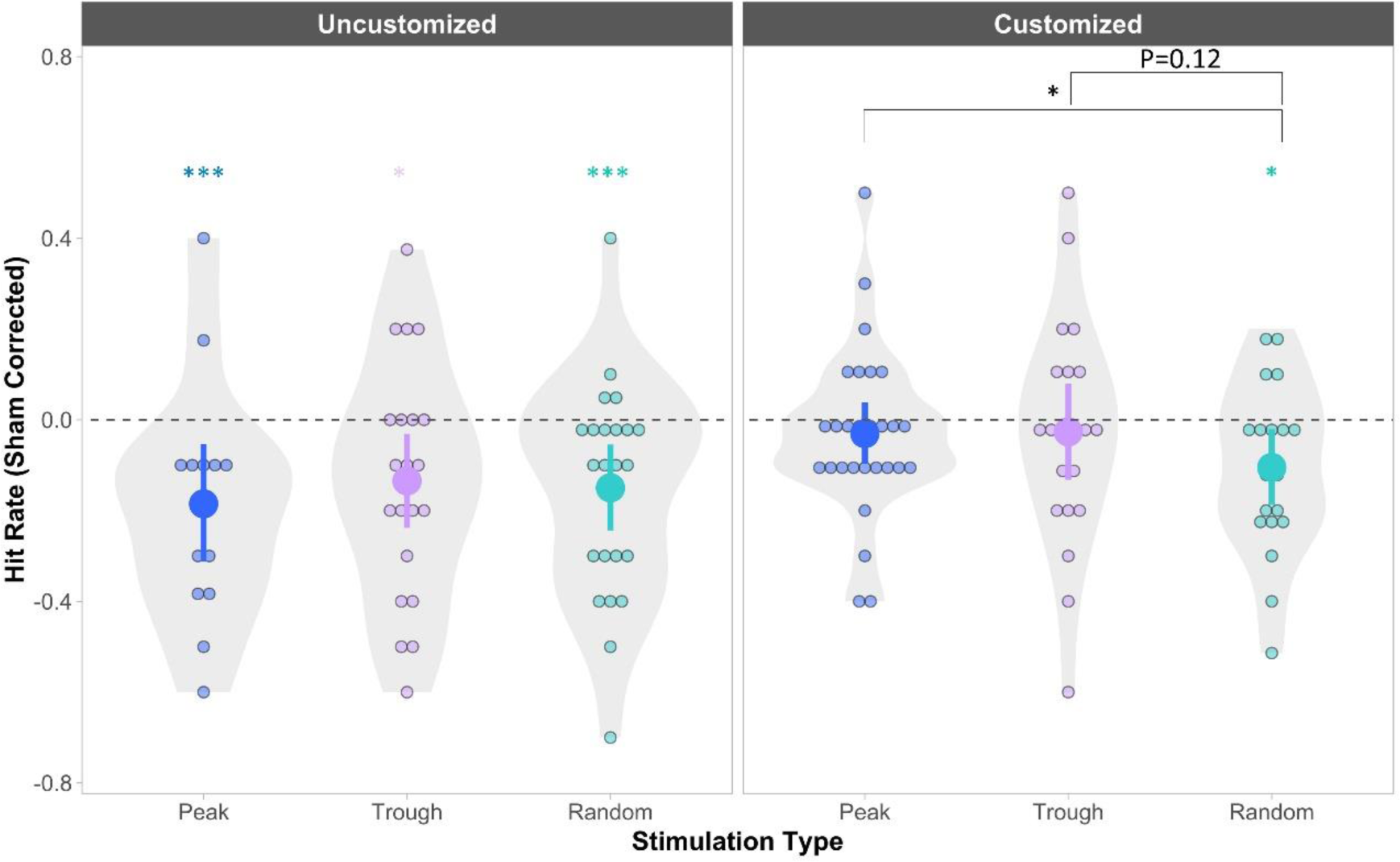
Model including Partially Customized sessions, labeling each peak and trough stimulation according to its respective customization. Average hit rate for *Non-Customized* (left) and Customized (right) sessions. There was a significant interaction between customization and stimulation type. Specifically, the impact of peak stimulation depended on whether its timing was customized (β = 0.93, SE = 0.35, *z* = 2.63, *p* = 0.008), and there was a similar trend for customization moderating the effect of trough stimulation (β = 079, SE = 0.45, *z*= 1.78, *p* = 0.08). Within the customized group, only random stimulation resulted in worse memory than sham (β = -0.49, SE = 0.24, *z* = -2.0, *p* = 0.045). Indeed, memory was superior in the peak compared to random stimulation conditions (β = -0.51, SE = 0.24, *z* = -2.17, *p* = 0.03). Sham is used as a baseline for the visualization of all stimulation types. Small dots indicate average rates per session; large dots indicate the group average and surrounding lines reflect the bootstrapped 95% confidence intervals. Note that data is averaged within sessions and sham corrected for ease of visualization, but reported statistics reflect interactions in the associated mixed model and follow-up contrasts performed with emmeans, which were estimated across trials. Colored significance markers denote differences from sham (baseline); markers contained in black bars mark differences between active stimulation conditions. * p < 0.05; ** p < 0.005; *** p < 0.0001

## Acknowledgements

I would like to acknowledge the summer work in the lab in helping with behavioural task Ryan Tian, analyzing some saccade metrics of previous iterations Alexandra Santos, and any lab members of Neuron to Brain who supported the work.

## Footnotes

1 We will be using orientation with recordings from the hippocampal fissure which is 180 degrees out of phase when recording from stratum radiatum in CA1 (Buzsaki et al., 1985)

## References

Andersson R, Larsson L, Holmqvist K, Stridh M, Nyström M (2017) One algorithm to rule them all? An evaluation and discussion of ten eye movement event-detection algorithms. Behav Res Methods 49:616–637.

Andrillon T, Nir Y, Cirelli C, Tononi G, Fried I (2015) Single-neuron activity and eye movements during human REM sleep and awake vision. Nat Commun 6:7884 Available at: http://www.nature.com/doifinder/10.1038/ncomms8884 [Accessed November 28, 2016].

Bates D, Mächler M, Bolker B, Walker S (2014) Fitting Linear Mixed-Effects Models using lme4. J Stat Softw Available at: http://arxiv.org/abs/1406.5823.

Bellana B, Mansour R, Ladyka-Wojcik N, Grady CL, Moscovitch M (2021) The influence of prior knowledge on the formation of detailed and durable memories. J Mem Lang 121:104264 Available at: 10.1016/j.jml.2021.104264.

Berman RA, Cavanaugh J, McAlonan K, Wurtz RH (2017) A circuit for saccadic suppression in the primate brain. J Neurophysiol 117:1720–1735 Available at: http://jn.physiology.org/lookup/doi/10.1152/jn.00679.2016.

Bush D, Bisby JA, Bird CM, Gollwitzer S, Rodionov R, Diehl B, McEvoy AW, Walker MC, Burgess N (2017) Human hippocampal theta power indicates movement onset and distance travelled. Proc Natl Acad Sci.

Buzsáki G (2002) Theta Oscillations in the Hippocampus. Neuron 33:325–340 Available at: http://www.cell.com/article/S089662730200586X/fulltext [Accessed December 21, 2014].

Buzsáki G (2005) Theta rhythm of navigation: Link between path integration and landmark navigation, episodic and semantic memory. Hippocampus 15:827–840.

Buzsaki G, Rappelsberger P, Kellenyi L (1985) Depth Profiles of Hippocampal Rhythmic Slow Activity (’Theta Rhythm’) Depend on Behaviour. Electroencephalograpt Clin Neurophysiol 61:77–88.

Chan JPK, Kamino D, Binns MA, Ryan JD (2011) Can changes in eye movement scanning alter the age-related deficit in recognition memory? Front Psychol 2 Available at: /pmc/articles/PMC3110339/ [Accessed February 28, 2021].

Coleshill SG, Binnie CD, Morris RG, Alarcón G, van Emde Boas W, Velis DN, Simmons A, Polkey CE, van Veelen CWM, van Rijen PC (2004) Material-Specific Recognition Memory Deficits Elicited by Unilateral Hippocampal Electrical Stimulation. J Neurosci 24.

Dolleman-Van Der Weel MJ, Lopes Da Silva FH, Witter MP (1997) Nucleus Reuniens Thalami Modulates Activity in Hippocampal Field CA1 through Excitatory and Inhibitory Mechanisms.

Eleore L, López-Ramos JC, Guerra-Narbona R, Delgado-García JM, Carlos Ló Pez-Ramos J, Guerra-Narbona R, Delgado-García JM (2011) Role of Reuniens Nucleus Projections to the Medial Prefrontal Cortex and to the Hippocampal Pyramidal CA1 Area in Associative Learning Izquierdo I, ed. PLoS One 6:e23538 Available at: www.plosone.org.

Enatsu R, Gonzalez-Martinez J, Bulacio J, Kubota Y, Mosher J, Burgess RC, Najm I, Nair DR (2015) Connections of the limbic network: a corticocortical evoked potentials study. Cortex 62:20–33 Available at: http://www.sciencedirect.com/science/article/pii/S0010945214002263 [Accessed March 4, 2016].

Estefan DP, Zucca R, Arsiwalla X, Principe A, Zhang H, Rocamora R, Axmacher N, Verschure PFMJ (2021) Volitional learning promotes theta phase coding in the human hippocampus. Proc Natl Acad Sci U S A 118:1–12.

Ezzyat Y et al. (2017) Direct Brain Stimulation Modulates Encoding States and Memory Performance in Humans. Curr Biol 27:1251–1258 Available at: 10.1016/j.cub.2017.03.028.

Ezzyat Y et al. (2018) Closed-loop stimulation of temporal cortex rescues functional networks and improves memory.

Fehlmann B, Coynel D, Schicktanz N, Milnik A, Gschwind L, Hofmann P, Papassotiropoulos A, de Quervain DJ-F (2020) Visual Exploration at Higher Fixation Frequency Increases Subsequent Memory Recall. Cereb Cortex Commun 1:1–14 Available at: https://academic.oup.com/cercorcomms/article/doi/10.1093/texcom/tgaa032/5874331 [Accessed April 16, 2021].

Gilboa A, Marlatte H (2017) Neurobiology of Schemas and Schema-Mediated Memory. Trends Cogn Sci 21:618–631 Available at: 10.1016/j.tics.2017.04.013.

Goyal A et al. (2018) Electrical stimulation in hippocampus and entorhinal cortex impairs spatial and temporal memory. J Neurosci 38:3049–17 Available at: http://www.jneurosci.org/lookup/doi/10.1523/JNEUROSCI.3049-17.2018.

Green JD, Arduini AA (1954) Hippocampal electrical activity in arousal. J Neurophysiol 17:533– 557.

Groppe DM, Bickel S, Dykstra AR, Wang X, Mégevand P, Mercier MR, Lado FA, Mehta AD, Honey CJ (2017) iELVis: An open source MATLAB toolbox for localizing and visualizing human intracranial electrode data. J Neurosci Methods 281:40–48 Available at: http://www.ncbi.nlm.nih.gov/pubmed/28192130 [Accessed August 25, 2017].

Halgren E, Wilson CL (1985) Recall deficits produced by afterdischarges in the human hippocampal formation and amygdala. Electroencephalogr Clin Neurophysiol 61:375–380.

Halgren E, Wilson CL, Stapleton JM (1985) Human medial temporal-lobe stimulation disrupts both formation and retrieval of recent memories. Brain Cogn 4:287–295.

Hampson RE, Song D, Robinson BS, Fetterhoff D, Dakos AS, Roeder BM, She X, Wicks RT, Witcher MR, Couture DE, Laxton AW, Munger-Clary H, Popli G, Sollman MJ, Whitlow CT, Marmarelis VZ, Berger TW, Deadwyler SA (2018) Developing a hippocampal neural prosthetic to facilitate human memory encoding and recall. J Neural Eng 15.

Hannula DE, Althoff RR, Warren DE, Riggs L, Cohen NJ, Ryan JD (2010) Worth a glance: using eye movements to investigate the cognitive neuroscience of memory. Front Hum Neurosci 4:166 Available at: http://www.ncbi.nlm.nih.gov/pubmed/21151363 [Accessed August 10, 2017].

Hasselmo ME, Bodelón C, Wyble BP (2002) A proposed function for hippocampal theta rhythm: separate phases of encoding and retrieval enhance reversal of prior learning. Neural Comput 14:793–817 Available at: http://www.mitpressjournals.org/doi/abs/10.1162/089976602317318965#.VrJZLTYrKRs [Accessed January 31, 2016].

Hasselmo ME, Stern CE (2014) Theta rhythm and the encoding and retrieval of space and time. Neuroimage 85:656–666.

Henderson JM, Williams CC, Falk RJ (2005) Eye movements are functional during face learning. Mem Cognit 33:98–106 Available at: http://www.springerlink.com/index/10.3758/BF03195300 [Accessed December 10, 2015].

Herweg NA, Solomon EA, Kahana MJ (2019) Theta Oscillations in Human Memory. Trends Cogn Sci:1–20 Available at: 10.1016/j.tics.2019.12.006.

Hoffman KL, Dragan MC, Leonard TK, Micheli C, Montefusco-Siegmund R, Valiante TA (2013) Saccades during visual exploration align hippocampal 3-8 Hz rhythms in human and non-human primates. Front Syst Neurosci 7:43 Available at: http://journal.frontiersin.org/article/10.3389/fnsys.2013.00043/abstract [Accessed September 14, 2015].

Hölscher C, Anwyl R, Rowan MJ (1997) Stimulation on the positive phase of hippocampal theta rhythm induces long-term potentiation that can Be depotentiated by stimulation on the negative phase in area CA1 in vivo. J Neurosci 17:6470–6477.

Huerta PT, Lisman JE (1995) Bidirectional synaptic plasticity induced by a single burst during cholinergic theta oscillation in CA1 in vitro. Neuron 15:1053–1063 Available at: http://www.sciencedirect.com/science/article/pii/0896627395900942 [Accessed December 21, 2015].

Huerta PT, Lisman JE (1996a) Low-frequency stimulation at the troughs of theta-oscillation induces long-term depression of previously potentiated CA1 synapses. J Neurophysiol 75:877–884.

Huerta PT, Lisman JE (1996b) Synaptic Plasticity During the Cholinergic Theta-Frequency Oscillation In Vitro. Hippocampus 61:58–61.

Hyman JM, Wyble BP, Goyal V, Rossi CA, Hasselmo ME (2003) Stimulation in hippocampal region CA1 in behaving rats yields long-term potentiation when delivered to the peak of theta and long-term depression when delivered to the trough. J Neurosci 23:11725–11731 Available at: http://www.jneurosci.org/content/23/37/11725.abstract [Accessed November 23, 2015].

Inman CS, Manns JR, Bijanki KR, Bass DI, Hamann S, Drane DL, Fasano RE, Kovach CK, Gross RE, Willie JT (2018) Direct electrical stimulation of the amygdala enhances declarative memory in humans. Proc Natl Acad Sci 115:98–103 Available at: http://www.pnas.org/lookup/doi/10.1073/pnas.1714058114.

Jacobs J et al. (2016) Direct Electrical Stimulation of the Human Entorhinal Region and Hippocampus Impairs Memory. Neuron 92:983–990 Available at: http://linkinghub.elsevier.com/retrieve/pii/S0896627316308364 [Accessed December 11, 2016].

Jutras MJ, Fries P, Buffalo EA (2013) Oscillatory activity in the monkey hippocampus during visual exploration and memory formation. Proc Natl Acad Sci U S A 110:13144–13149 Available at: http://www.pnas.org/cgi/doi/10.1073/pnas.1302351110 [Accessed October 9, 2015].

Kafkas A, Montaldi D (2011) Recognition memory strength is predicted by pupillary responses at encoding while fixation patterns distinguish recollection from familiarity. Q J Exp Psychol 64:1971–1989.

Katz CN, Patel K, Talakoub O, Groppe D, Hoffman KL, Valiante TA (2020) Differential Generation of Saccade, Fixation, and Image-Onset Event-Related Potentials in the Human Mesial Temporal Lobe. Cereb Cortex 30:5502–5516.

Katz CN, Schjetnan AG, Patel K, Barkley V, Hoffman KL, Kalia SK, Duncan KD, Valiante TA (2022) A corollary discharge mediates saccade-related inhibition of single units in mnemonic structures of the human brain. Curr Biol Available at: 10.1016/j.cub.2022.06.015 [Accessed July 6, 2022].

Keller CJ, Honey CJ, Mégevand P, Entz L, Ulbert I, Mehta AD (2014) Mapping human brain networks with cortico-cortical evoked potentials. Philos Trans R Soc Lond B Biol Sci 369 Available at: http://www.pubmedcentral.nih.gov/articlerender.fcgi?artid=4150303&tool=pmcentrez&rendertype=abstract [Accessed February 12, 2016].

Kerrén C, Linde-Domingo J, Hanslmayr S, Wimber M (2018) An Optimal Oscillatory Phase for Pattern Reactivation during Memory Retrieval. Curr Biol 28:3383–3392.e6 Available at: http://www.ncbi.nlm.nih.gov/pubmed/30344116.

Kocabicak E, Temel Y, Höllig A, Falkenburger B, Tan SK (2015) Current perspectives on deep brain stimulation for severe neurological and psychiatric disorders. Neuropsychiatr Dis Treat 11:1051–1066 Available at: http://www.pubmedcentral.nih.gov/articlerender.fcgi?artid=4399519&tool=pmcentrez&rendertype=abstract [Accessed February 5, 2016].

Kragel JE, VanHaerents S, Templer JW, Schuele S, Rosenow JM, Nilakantan AS, Bridge DJ (2020) Hippocampal theta coordinates memory processing during visual exploration. Elife 9 Available at: https://elifesciences.org/articles/52108.

Kragel JE, Voss JL (2022) Looking for the neural basis of memory. Trends Cogn Sci 26:53–65.

Krout KE, Loewy AD, Westby GW, Redgrave P (2001) Superior colliculus projections to midline and intralaminar thalamic nuclei of the rat. J Comp Neurol 431:198–216.

Kubota Y, Enatsu R, Gonzalez-Martinez J, Bulacio J, Mosher J, Burgess RC, Nair DR (2013) In vivo human hippocampal cingulate connectivity: a corticocortical evoked potentials (CCEPs) study. Clin Neurophysiol 124:1547–1556 Available at: http://www.sciencedirect.com/science/article/pii/S1388245713001004 [Accessed February 12, 2016].

Kucewicz MT et al. (2018a) Evidence for verbal memory enhancement with electrical brain stimulation in the lateral temporal cortex. Brain 141:971–978.

Kucewicz MT, Berry BM, Kremen V, Miller LR, Ezzyat Y, Stein JM, Wanda P, Sperling MR, Davis KA, Jobst BC, Gross RE, Lega B, Matt S, Rizzuto DS, Kahana MJ, Worrell GA (2018b) Electrical Stimulation Modulates High Activity and Human Memory Performance. 5:1–14.

Lacruz ME, Valentín A, Seoane JJG, Morris RG, Selway RP, Alarcón G (2010) Single pulse electrical stimulation of the hippocampus is sufficient to impair human episodic memory. Neuroscience 170:623–632 Available at: http://www.sciencedirect.com/science/article/pii/S0306452210008894 [Accessed October 9, 2015].

Larson J, Wong D, Lynch G, Larson (1986) Patterned stimulation at the theta frequency is optimal for the induction of hippocampal long-term potentiation. Brain Res 368:347–350.

Law CSH, Stan Leung L (2018) Long-term potentiation and excitability in the hippocampus are modulated differently by θ rhythm. eNeuro 5.

Leão R, Mikulovic S, Leão KE, H M, H G, A E, K P, A E, LM L, AB T, K K (2012) OLM interneurons differentially modulate CA3 and entorhinal inputs to hippocampal CA1 neurons. Nat Neurosci 15:1524–1530.

Lin CJ et al. (2014) Developing and Evaluating a Target-Background Similarity Metric for Camouflage Detection Osorio D, ed. PLoS One 9:e87310 Available at: http://dx.plos.org/10.1371/journal.pone.0087310 [Accessed November 18, 2016].

Lowet E, Kondabolu K, Zhou S, Mount RA, Wang Y, Ravasio CR, Han X (2022) Deep brain stimulation creates informational lesion through membrane depolarization in mouse hippocampus. Nat Commun 13:1–15.

Lurie SM, Kragel JE, Schuele SU, Voss JL (2022) Human hippocampal responses to network stimulation vary with theta phase Abbreviated title: Theta phase gates hippocampal responsivity. :1–26 Available at: 10.1101/2022.02.28.482345.

M. Aghajan Z, Schuette P, Fields TA, Tran ME, Siddiqui SM, Hasulak NR, Tcheng TK, Eliashiv D, Mankin EA, Stern J, Fried I, Suthana N (2017) Theta Oscillations in the Human Medial Temporal Lobe during Real-World Ambulatory Movement. Curr Biol 27:3743–3751.e3 Available at: 10.1016/j.cub.2017.10.062.

Mankin EA, Aghajan ZM, Schuette P, Tran ME, Tchemodanov N, Titiz A, Kalender G, Eliashiv D, Stern J, Weiss SA, Kirsch D, Knowlton B, Fried I, Suthana N (2021) Stimulation of the right entorhinal white matter enhances visual memory encoding in humans. Brain Stimul 14:131–140 Available at: 10.1016/j.brs.2020.11.015.

Mao D, Avila E, Caziot B, Laurens J, Dickman JD, Angelaki DE (2021) Spatial modulation of hippocampal activity in freely moving macaques. Neuron Available at: https://linkinghub.elsevier.com/retrieve/pii/S0896627321007054 [Accessed October 15, 2021].

Maris E, Oostenveld R (2007) Nonparametric statistical testing of EEG-and MEG-data. J Neurosci Methods 164:177–190.

McCartney H, Johnson AD, Weil ZM, Givens B (2004) Theta reset produces optimal conditions for long-term potentiation. Hippocampus 14:684–687 Available at: http://www.ncbi.nlm.nih.gov/pubmed/15318327.

Meister MLRR, Buffalo EA (2016) Getting directions from the hippocampus: The neural connection between looking and memory. Neurobiol Learn Mem 134:135–144 Available at: http://www.sciencedirect.com/science/article/pii/S1074742715002348 [Accessed December 30, 2015].

Merkow MB, Burke JF, Ramayya AG, Sharan AD, Sperling MR, Kahana MJ (2016) Stimulation of the human medial temporal lobe between learning and recall selectively enhances forgetting. Brain Stimul:1–6 Available at: http://linkinghub.elsevier.com/retrieve/pii/S1935861X1630393X.

Miller JP, Sweet JA, Bailey CM, Munyon CN, Luders HO, Fastenau PS (2015) Visual-spatial memory may be enhanced with theta burst deep brain stimulation of the fornix: A preliminary investigation with four cases. Brain 138:1833–1842 Available at: http://www.ncbi.nlm.nih.gov/pubmed/26106097 [Accessed September 14, 2016].

Molitor RJ, Ko PC, Hussey EP, Ally BA (2014) Memory-related eye movements challenge behavioral measures of pattern completion and pattern separation. Hippocampus 24:666–672.

Morales GJ, Ramcharan EJ, Sundararaman N, Morgera SD, Vertes RP (2007) Analysis of the Actions of Nucleus Reuniens and the Entorhinal Cortex on EEG and Evoked Population Behavior of the Hippocampus. In: 2007 29th Annual International Conference of the IEEE Engineering in Medicine and Biology Society, pp 2480–2484. IEEE.

Nokia MS, Waselius T, Mikkonen JE, Wikgren J, Penttonen M (2015) Phase matters: responding to and learning about peripheral stimuli depends on hippocampal θ phase at stimulus onset. Learn Mem 22:307–317 Available at: http://learnmem.cshlp.org/content/22/6/307.abstract.

Olsen RK, Sebanayagam V, Lee Y, Moscovitch M, Grady CL, Rosenbaum RS, Ryan JD (2016) The relationship between eye movements and subsequent recognition: Evidence from individual differences and amnesia. Cortex 85:182–193.

Ryan JD, Shen K, Liu ZX (2020) The intersection between the oculomotor and hippocampal memory systems: empirical developments and clinical implications. Ann N Y Acad Sci 1464:115–141.

Sakalar E, Klausberger T, Lasztóczi B (2022) Neurogliaform cells dynamically decouple neuronal synchrony between brain areas. Science (80-) 377:324–328.

Scheel N, Wulff P, de Mooij-van Malsen JG (2020) Afferent connections of the thalamic nucleus reuniens in the mouse. J Comp Neurol 528:1189–1202.

Siegle JH, Wilson MA (2014) Enhancement of encoding and retrieval functions through theta phase-specific manipulation of hippocampus. Elife 2014:1–18.

Sobotka S, Ringo JL (1997) Saccadic eye movements, even in darkness, generate event-related potentials recorded in medial sputum and medial temporal cortex. Brain Res 756:168–173.

Sobotka S, Zuo W, Ringo JL (2002) Is the functional connectivity within temporal lobe influenced by saccadic eye movements? J Neurophysiol 88:1675–1684 Available at: http://jn.physiology.org/content/88/4/1675.long [Accessed June 13, 2017].

Sommer MA, Wurtz RH (2002) A pathway in primate brain for internal monitoring of movements. Science 296:1480–1482.

Stewart M, Fox SE (1991) Hippocampal theta activity in monkeys. Brain Res 538:59–63.

Suthana N, Aghajan ZM, Mankin EA, Lin A (2018) Reporting Guidelines and Issues to Consider for Using Intracranial Brain Stimulation in Studies of Human Declarative Memory. Front Neurosci 12.

Talakoub O, Gomez Palacio Schjetnan A, Valiante TA, Popovic MR, Hoffman KL (2016) Closed-Loop Interruption of Hippocampal Ripples through Fornix Stimulation in the Non-Human Primate. Brain Stimul 9:911–918 Available at: http://linkinghub.elsevier.com/retrieve/pii/S1935861X16302029 [Accessed August 7, 2017].

Titiz AS, Hill MRH, Mankin EA, M Aghajan Z, Eliashiv D, Tchemodanov N, Maoz U, Stern J, Tran ME, Schuette P, Behnke E, Suthana NA, Fried I (2017) Theta-burst microstimulation in the human entorhinal area improves memory specificity. Elife 6:e29515 Available at: https://elifesciences.org/articles/29515.

Wang Z, Bovik A (2002) A universal image quality index. IEEE Signal Process Lett 9:81–84.

Wang Z, Bovik AC, Sheikh HR, Simoncelli EP (2004) Image quality assessment: From error visibility to structural similarity. IEEE Trans Image Process 13:600–612.

Womelsdorf T, Valiante TA, Sahin NT, Miller KJ, Tiesinga P (2014) Dynamic circuit motifs underlying rhythmic gain control, gating and integration. Nat Neurosci 17:1031–1039 Available at: http://www.ncbi.nlm.nih.gov/pubmed/25065440.

Zhou B, Lapedriza A, Khosla A, Oliva A, Torralba A (2018) Places: A 10 Million Image Database for Scene Recognition. IEEE Trans Pattern Anal Mach Intell 40:1452–1464.

